# NucleoSeeker: Precision Filtering of RNA Databases to Curate High-Quality Datasets

**DOI:** 10.1101/2024.12.06.626307

**Authors:** Utkarsh Upadhyay, Fabrizio Pucci, Julian Herold, Alexander Schug

## Abstract

The structural prediction of biomolecules via computational methods complements the often involved wet-lab experiments. Unlike protein structure prediction, RNA structure prediction remains a significant challenge in bioinformatics, primarily due to the scarcity of annotated RNA structure data and its varying quality. Many methods have used this limited data to train deep learning models but redundancy, data leakage and bad data quality hampers their performance. In this work, we present NucleoSeeker, a tool designed to curate high-quality, tailored datasets from the Protein Data Bank (PDB) database. It is a unified framework that combines multiple tools and streamlines an otherwise complicated process of data curation. It offers multiple filters at structure, sequence and annotation levels, giving researchers full control over data curation. Further, we present several use cases. In particular, we demonstrate how NucleoSeeker allows the creation of a non-redundant RNA structure dataset to assess AlphaFold3’s performance for RNA structure prediction. This demonstrates NucleoSeeker’s effectiveness in curating valuable non-redundant tailored datasets to both train novel and judge existing methods. NucleoSeeker is very easy to use, highly flexible and can significantly increase the quality of RNA structure datasets.

## 1 Introduction

Deep learning (DL) technology has given a significant boost to scientific research by providing powerful tools for data analysis, pattern recognition and prediction [1]. It strongly impacted the computational structural biology community enabling the recent breakthroughs such as AlphaFold [2] providing a massive improvement in both speed and accuracy for protein structure prediction.

Prior methods based on statistical inference such as direct coupling analysis (DCA) [3] gave a glimpse at the value hidden within the evolution of biomolecular sequences, enabling the statistical inference of spatial adjacencies to guide structure prediction tools [4, 5]. Transformer networks as used by models such as AlphaFold [2] leverage the information of protein evolution as found in sequence data to derive structural information. While these approaches are highly successful [6], they require abundant training data.

Thus, scarcity of data prohibits the direct transfer of these methods to other biomolecules, e.g. RNA. Moreover, the available RNA structural data suffers not only from its limited size but also from high redundancy and low data quality. Specifically, the current version of the Protein Data Bank (PDB) [7] contains a large number of highly similar RNA structures, structures with poor resolution, a significant number of hybrids (Protein/RNA, DNA/RNA and others) and a considerable proportion of very short sequences (fewer than twenty residues) (see Fig. 1a and Figure 1 in Supplementary Information).

**Figure 1.**
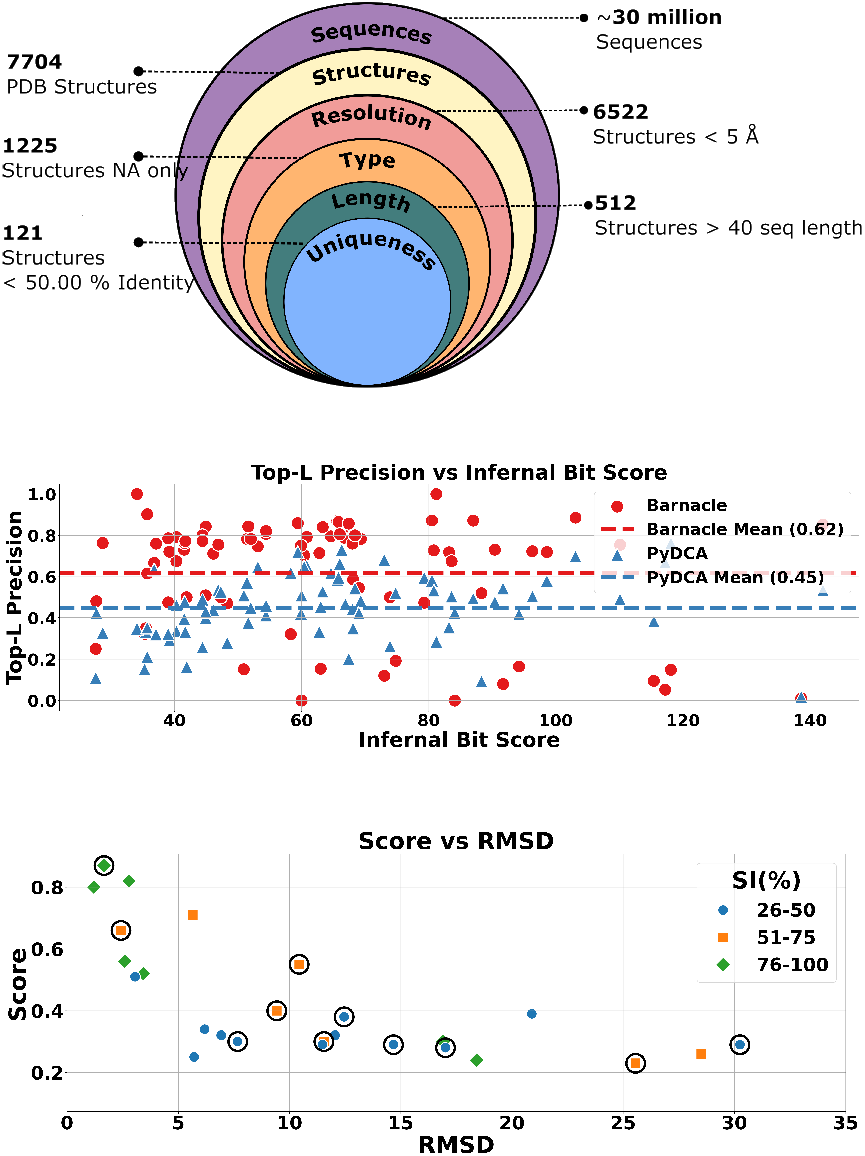
(a, top) **Hierarchical classification of RNA data**: from 30 million sequences to 121 unique structures. This nested diagram illustrates the progressive filtering of RNA information, showcasing the rarity of well-characterized, unique structures among the vast sea of known sequences. Each layer represents increasingly stringent criteria. (b, middle) **Barnacle and PyDCA contact prediction performance**: Top-L precision of two RNA contact prediction tools, Barnacle (red circles) and PyDCA (blue triangles), for the 𝒟_𝒞_ structures as a function of their Infernal Bit Score with the corresponding RFAM families (x-axis). (c, bottom) **AlphaFold pTM score vs RMSD**: AlphaFold pTM score as a function of the Root Mean Square Deviation (RMSD) between AlphaFold3 predictions and experimental structures in Å, categorized by sequence identity (SI%) levels.

Such properties often cause DL models to overfit and generalize poorly. Assessing DL models is also challenging because data leakage, stemming from improper splits between training and test sets, can lead to an overestimation of the model’s performance [8, 9].

Furthermore, reproducibility also becomes impaired. Experiments like RNA-Puzzles [10] and CASP [11], where computational algorithms have to blindly predict RNA structures, provide a valid evaluation method. However, results [11] suggest that due to the above-mentioned problems, DL models currently perform worse than physics-based approaches in RNA structure prediction task.

The development of curated datasets for training and testing models is essential to addressing these issues. For instance, following strict filtering processes and manual curation as in [12, 13] can be highly effective. Here, we introduce NucleoSeeker, an easy-to-use software that provides extensive flexibility and control for curating RNA datasets from structures deposited in the PDB database in a fully automated manner.

## 2 Methods

NucleoSekeer is a python library that can be directly used as a command-line tool with limited dependencies (i.e. Biopython, Pandas, Numpy and Requests). It handles downloading and applying filters to create a dataset.

### 2.1 Dataset Access

Initially, no structures are downloaded from the RCSB PDB database. Instead, we first use the `Search API` of the PDB to retrieve all IDs for a specified structure determination method and a given polymer entry type. These IDs are then processed through a `GraphQL` query, which fetches predefined attributes, such as the experimental method used, resolution and many more for each structure (see section 1 of Supplementary Information for details). This API-based approach ensures that our tool generates the most up-to-date dataset without requiring any code modifications. This module yields a data frame `DF` with all requested IDs and their corresponding attributes.

To refine the dataset, we use three different kinds of filters in our software that allow the users to specify their requirements for various levels from the individual chain to multiple structures; all the filter modules in the package are also available as standalone modules.

### 2.2 Dataset Creator

The results of the filtering operations are combined to generate a dataset of RNA structure. Infernal [14] is then used to search for RFAM families [15] of the filtered structures. Users can specify an E-value threshold for the RNA family hits that specifies the statistical significance of the result (refer to the user guide of Infernal [14] for more details). A lower E-value indicates a more significant result, effectively controlling the strictness of the family search.

The output of NucleoSeeker consists of a list of RNA chains along with corresponding information, such as RFAM classification, PDB code and the corresponding number of chains, structure resolution, experimental method used, and year of release. Note that, in the case of complexes, only the RNA chains that meet all the specified criteria are selected.

### 2.3 Filtering Mechanisms

In our filtering approach, we follow the hierarchy used by the RCSB PDB database to organize structures. More in detail, we use the three common levels `ENTRY, ENTITY` and `INSTANCE`, which form the basis of the different modules.

Here, we will briefly describe the functionality of the different modules and how they integrate to generate the final dataset. An exhaustive list of arguments and parameters for each module can be found in the Supplementary Information. Note, that the word `Polymer` is used in multiple parameters across modules, and each of them carries a different meaning (see Supplementary Information).

### Metadata

We use this module to apply filters based on the metadata stored in the `DF`. This offers a comprehensive set of filters for structural attributes, e.g. users can filter structures based on the experimental methods used for determination, such as X-ray diffraction, NMR, or Cryo-EM, ensuring that only structures determined by these techniques are included. The module also offers a resolution threshold filter, allowing users to exclude structures with resolutions outside a specified value. Similarly, structures can be filtered by their release year, allowing the inclusion of only those resolved before a specific year or within a particular range of years. Furthermore, the module supports selection based on polymer entity types, enabling users to customize their datasets by focusing on specific polymers or excluding polymer complexes. Additionally, the keyword filter provides the ability to include or exclude structures based on specific terms, offering further refinement of the dataset to meet users’ requirements.

### Individual Structure

The `IndividualStructure` module analyzes the composition of structures for further processing, i.e. it parses individual files. It examines each structure’s polymer type, nucleotide or residue integrity, and the length of each chain. This module can function independently to download and verify PDB files based on specified criteria when given a PDB ID. Utilizing a PDB parser, it extracts and analyzes structural information, allowing users to specify parameters such as the desired polymer instance type and sequence length to filter out chains that do not meet these criteria. This module enables the easy removal of short sequences and the extraction of chains containing only RNA.

### Structure Comparison

This filter level combines our filtered DF from the `Metadata` filter and the `IndividualStructure` filter by applying the latter to each structure in the `DF`. We also integrate sequence alignment tools, such as Clustal Omega [16] and Emboss [17], to filter structures based on sequence identity (SI). If the SI between two structures exceeds a certain threshold, We select the structure with the highest resolution to minimize redundancy in the dataset while ensuring good overall structure resolution.

A sequence identity matrix is created using the specified alignment tools. However, since `EMBOSS` processes sequences in pairs, impacting performance, it is generally recommended to use `Clustal Omega` for most applications. These filters, whether used individually or collectively, provide an unprecedented level of control and flexibility in curating datasets from the PDB database. Consequently, researchers can create more targeted and refined datasets, significantly enhancing the accuracy and reliability of their RNA structure prediction protocols.

## 3 Results

Well-curated and non-redundant datasets serve two important goals: they improve training outcomes and are crucial for assessing method performance. Here, we show that NucleoSeeker can be used to attain this goal. In particular, we highlight two examples where our tool can create datasets for assessing RNA contact prediction methods and evaluating AlphaFold3 [18] performance on RNA structure prediction. The manual curation of such datasets is time-consuming and error-prone and can lead to non-systematic biases. We show the ease of curating such datasets using NucleoSeeker and believe it can be used to prepare datasets for machine learning algorithms in a similar way.

### 3.1 Use-case 1: Automated RNA structures curation for assessing contact prediction

We used NucleoSeeker to create a well-curated and non-redundant dataset starting from all the 7704 RNA structures available in the PDB database (accessed in July 2024). We selected only RNA structures resolved by X-ray crystallography, with a resolution below 3.6*Å* and a maximum pair-wise sequence identity of 50%. Since we wanted to create a dataset of RNA-only structures we used `‘pdbx keywords’=‘RNA’`. We ended up with 117 structures, out of which 88 have an associated RFAM family [15]. These parameters are the same as those used in the construction of the dataset curated in [12], in which only 69 families were included (PDB database accessed in 2020). We note that although there has been an increase in the number of resolved RNA structures over the years, the data remains scarce and significant improvements are needed to train DL models on these limited data.

This dataset labelled with 𝒟_𝒞_ is then used to assess the performance of two unsupervised RNA contact prediction methods, namely PyDCA [19] and Barnacle [20]. In Fig. 1.b, we present the performance of these two methods on 𝒟_𝒞_, as measured by the Precision at rank L, which represents the proportion of correctly predicted nucleotide contacts among the top L predictions.

We note that the two methods reach good performances with Precision equal to about 0.62 for Barnacle and 0.45 for PyDCA when averaged on all structures belonging to 𝒟_𝒞_. Barnacle [20], which utilizes data-efficient machine learning, generally achieves higher precision than PyDCA, which relies on the pseudo-likelihood maximization direct coupling analysis approach as it is more effective in leveraging MSA information to predict contacts. Table 2 of Supplementary Information contains all the top L precision values.

There is no clear trend between bit score and precision, as both tools demonstrate variability in performance across different bit scores. This is because not only the bit score but also the effective number of sequences in the RFAM family plays a significant role in the ability of methods to extract structural information from MSA [12].

### 3.2 Use-case 2: Automated RNA structures curation for quick assessment of AlphaFold3 capabilities on RNA

AlphaFold3 [18] shows remarkable promise in protein structure prediction and also promises to predict RNA and RNA complexes. Here, we assess its capabilities to predict the structure of unseen RNA sequences. To conduct such a comprehensive and unbiased assessment, we meticulously crafted two datasets using NucleoSeeker.

Our first dataset, which we designated 𝒟_22_, comprised 213 RNA structures solved before 2023 as the training dataset used by AlphaFold3 was derived from the PDB accessed on January 2023. We applied stringent selection criteria to ensure the highest quality and representativeness of this dataset (for a detailed description of these criteria see Supplementary Information Section 3.2).

Complementing this, we created a second dataset, 𝒟_23−24_, consisting of 27 structures solved in 2023 and 2024. This more recent dataset was important for assessing Alphafold3’s performance on newly determined structures. Since we know, that sequence identity is an important factor in reducing redundancy, we categorize the structures in 𝒟_23−24_ based on their similarity to those in 𝒟_22_ by calculating pairwise sequence identity (SI) between all structures across both datasets and dividing the 𝒟_23−24_ entries in three subclasses according to their SI: below 50%, between 51% and 75% and above 76% (see Supplementary Table 3).

In Fig. 1.c, we plot the pTM score, a confidence metric for the predicted structure from AlphaFold3 [18], against the RMSD between the predicted and experimental structure, for all structures in 𝒟_23−24_ and according to their sequence identity with the closest match in the AF3 training se. Our quick assessment revealed that the AlphaFold3 confidence score genuinely provides a good indication of prediction quality, as high pTM scores correspond to RMSD values generally smaller than 5 Å. Additionally, the performance of AF3 improves for structures with higher sequence identity, as indicated by the green diamonds. This observation suggests that the model’s accuracy is influenced by the similarity between the target structure and those in AF3 training data (check section 3.2 of the Supplementary Information for more details).

While Alphafold3 shows promising performance, especially for RNAs with some degree of similarity to known structures, this preliminary analysis shows that there is still room for improvement in predicting novel or highly divergent RNA structures. As we move forward, these insights will guide our efforts to refine and enhance RNA structure prediction methodologies.

## 4 Discussion

NucleoSeeker targets to complement the development of efficient deep learning methods for RNA structure prediction, by providing a robust and flexible method for curating high-quality datasets from the PDB database. One of the key strengths of this software is its ability to apply a wide range of filters at both the structure and sequence levels, allowing researchers to create highly specific and relevant datasets tailored to their particular research needs. This functionality is particularly valuable given the challenges associated with RNA data, such as high redundancy, poor resolution, and the presence of hybrid structures. Moreover, the system is designed to ensure that datasets remain up-to-date, even in the rapidly evolving field of RNA research, where new structures are continually being determined and added to the database. Additionally, the modular design of NucleoSeeker allows its components to be used independently or in combination, providing researchers with a high degree of flexibility. Whether the goal is to filter specific structures, analyze polymer chains, or reduce dataset redundancy, it offers the tools needed to achieve these objectives efficiently in a simple way.

## 5 Data Availability

The code for NucleoSeeker is available on GitHub at https://github.com/theuutkarsh/nucleoseeker and on Zenodo https://doi.org/10.5281/zenodo.13843170.

The package is also available through PyPi, Python Package Index.

## 6 Acknowledgments

The authors gratefully acknowledge the Gauss Centre for Supercomputing e.V. (www.gauss-centre.eu) for funding this project by providing computing time through the John von Neumann Institute for Computing (NIC) on the GCS Supercomputer JUWELS at Jülich Supercomputing Centre (JSC) [21]. The authors gratefully acknowledge computing time on the supercomputer JURECA [22] at Forschungszentrum Jülich under the grant name EatsRNA.

