## Supplementary Information for "NucleoSeeker: Precision Filtering of RNA Databases to Curate High-Quality Datasets"

December 2, 2024

**4 Supplementary Tables** **7**

**5 Supplementary Figures** **9**

### 1 Background

Deep learning (DL) has accelerated scientific research, leading to breakthroughs like AlphaFold's accurate protein structure prediction. Historically, direct coupling analysis (DCA) leveraged crucial insights from biomolecular sequence evolution to aid in structure prediction. Transformer networks have further imporved the accuracy of linking protein evolution to structural information, but these models need abundant data.

Applying these methods to RNA is challenging due to scarce data and other constraints (see figure 1). The Protein Data Bank (PDB) contains many similar RNA structures, low-resolution data, hybrids, and short sequences, leading to overfitting and poor generalization in DL models. Evaluating these models is difficult due to potential data leakage and impaired reproducibility.

Developing highly curated datasets is essential to address these issues and improve RNA structure prediction. Thus, we developed NucleoSeeker, which filters RNA structures from the PDB database in an automated manner. NucleoSeeker consists of multiple modules, which can be utilized individually or consecutively as a pipeline handling all required steps from downloading the specified data, to filtering the structures and creating the final dataset. Here we describe each of the modules and related use cases.

First, before any filters are used we extract metadata of structures in a tabular format (a data frame `DF`. We use the following attributes:-

- `rcsb_id`
- `exptl_method`
- `release_date`
- `polymer_entity_instance_count`
- `polymer_entity_count_RNA`
- `resolution`
- `selected_polymer_entity_types`
- `pdbx_keywords`

These attributes were chosen because they contain significant data about each structure and facilitate efficient filtering based on specific criteria. For example, we can filter structures based on attributes such as the number of chains and types of polymers. This approach optimizes performance by initially filtering structures based on attributes, thereby reducing the number of structure files to be downloaded. Subsequently, we analyze a smaller, refined set of structure files.

### 2 Modules

#### 2.1 Metadata Filter

We use this module to apply metadata filters on the data stored in the DF. Since we don't analyse PDB structures directly, we use this module to filter our DF as per requirements. It has filters corresponding to each attribute of the structures in the DF.

1. `exptl_method`: Users can specify experimental techniques used for structure determination such as X-ray diffraction, NMR, Cryo-EM etc.
2. `resolution`: The user can choose a resolution threshold and the command provides all structures with a resolution below that value.
3. Users can specify a year, and all structures resolved after that year are filtered out. Moreover, a `year_range` is also accepted to provide structures resolved only in those years.
4. The `selected_polymer_entity_types` filter is another valuable feature of our tool. It provides options to select the type of polymers in our structure. Whether researchers are interested in RNA, DNA, or protein structures, this filter enables them to tailor their dataset accordingly.
5. The `polymer_entity_instance_count` filter allows the user to filter the dataset based on the number of chains of a given polymer type in the structure.
6. The `polymer_entity_count_RNA` filter allows to set the precise number of RNA chains wanted in the dataset.
7. Lastly, the `pdbx_keywords` filter enables users to include or exclude structures based on specific keywords.

#### 2.2 Individual Structure Filter

The next module of our tool checks individual PDB files at a more detailed level. Like all other modules, this also works standalone, for a specific PDB ID it can download and check if it satisfies the given criteria.

It takes the following parameters -

1. `pdb_parser`: This parameter accepts an instance of a PDB parser. It is used to parse the structure and extract necessary information for further filtering.
2. `pdb_id`: This is where you need to provide the PDB ID of the structure you want to filter.
3. `polymer_type`: This parameter accepts the type of polymer you want to keep in your dataset.

4. `sequence_length`: This parameter allows you to specify the length of the sequence. Polymer chains with sequence lengths greater than this value are rejected.

5. `auto_download`: This parameter when set to True; allows automatic download of the PDB file if it is not already present in the local directory. If `auto_download` is set to False, it will throw an error when the PDB file is not found locally.

For a given PDB ID this module returns which chain, along with its sequence, in the structure falls under the requested parameters.

#### 2.3 Structure Comparison Filter

This module takes the following parameters, a few of which are the same as the IndividualStructureFilter-

1. `polymer_type`: This parameter determines the type of biopolymer to be included in the dataset. This flexibility allows users to tailor their datasets according to specific research needs, whether they're primarily interested in RNA, DNA, or protein structures.

2. `sequence_length`: This parameter sets a threshold for the length of the sequences to be included in the dataset. Any polymer chains with sequence lengths exceeding this value will be excluded. This feature helps to maintain a manageable dataset size and ensures the focus on sequences of interest.

3. `sequence_identity`: This parameter is crucial for reducing redundancy in the dataset. It sets a cutoff value for sequence identity. If two structures have a sequence identity greater than this value, the structure with the higher resolution is selected, ensuring a diverse yet high-quality dataset.

4. `auto_download`: This parameter controls whether the PDB file for a given structure is automatically downloaded if it's not already present in the local directory. If set to False, the absence of the required PDB file in the local directory will trigger an error.

5. `alignment_tool`: This parameter allows users to select the sequence alignment tool of their choice between ClustalOmega and Emboss. These tools are integrated into the software, to calculate sequence identity between given PDB files. This feature provides flexibility according to user preference and the specific requirements of different research projects. We recommend using ClustalOmega for better performance.

### 3 Usage Examples

To generate a new dataset, the first step is to setup NucleoSeeker by following the instructions given here here. After that, you can write the following command in your terminal and generate a dataset.

```
export DATA_PATH=/path/to/data # the dataset is saved here
nucleoseeker \
    --dataset_name test_dataset \
    --rfam_cm_path your/rfam/path \
    --exptl_method "X-RAY DIFFRACTION" \
    --resolution 3.6 \
    --year_range 2024 \
    --save 1 \
    --dend 500
```

After using this command a directory with the name `test_dataset` will be created in the `DATA_PATH` directory with the following subdirectories:

```
|--DATA_PATH
   |--pdb_files
   |--test_dataset
      |--files
      |--sequences
      |--clean_tblout.tblout
      |--cmscan.out
      |--combined.fasta
      |--fam_pdb_chain.csv
      |--final.fasta
      |--raw_experimental_RNA_0_500.csv
      |--sequence_identity_mat_clustal.csv
      |--tblout.tblout
```

- This tool generates various files at each level of filter. The first file that is generated is the `raw*.csv` file which contains the raw data from the PDB database.
- Then the `combined.fasta` file is generated which contains sequences used in sequence identity calculation by `Clustal Omega` and `Emboss`. This file is obtained after applying `StructureLevelFilter` and `PDBFilter` on the raw data.
- The `sequence_identity_mat_clustal(/emboss).csv` file contains the sequence identity matrix obtained from `Clustal Omega` or `Emboss`.
- The `final.fasta` file contains the final sequences in fasta format, these are the final sequences and if you don't want to analyse families then this is the final output.
- After this `cmscan.out`, `tblout.tblout`, `clean_tblout.tblout` files are generated which are the output of `Infernal` tool. The `fam_pdb_chain.csv` file contains the mapping of the family and the PDB chain.

- The fam_pdb_chain.csv is obtained after family search by Infernal tool. This is your final output if you want to analyse families.
- The files directory contains the dataframe and list for structures at each level of filter. These are generated only when save=1 is used.
- The sequences directory sequences for all the final structures in individual fasta files.

#### 3.1 Contact Maps Analysis: PyDCA vs. Barnacle

We generated a dataset for analysing the performance of PyDCA and Barnacle, using the following command: Command Used:

```
export DATA_PATH=/path/to/data # the dataset is saved here
nucleoseeker \
    --dataset_name name \
    --rfam_cm_path your/rfam/path \
    --exptl_method "X-RAY DIFFRACTION" \
    --resolution 3.6 \
    --year_range 2024 \
    --pdbx_keywords "RNA" \
    --save 1
```

Both of these software generate contact maps thus to quantify their performance using top-L precision we generated target contact maps. We define contact when the distance between the heavy atoms of two residues is less than 8 Å, and the residues are separated by 4 residues in between. The structure 1duh had only one contact leading to no result, we have represented it by nan, see supplementary table 2. Figure 3.1 shows the top-L values for each structure in the dataset.

#### 3.2 Alphafold3 Analysis

For analysing the performance of Alphafold3 we generated two different datasets. The first was up to 2022 ($\mathcal{D}_{22}$), meaning structures in this dataset were likely used for training, while the second was from 2023 to June 2024 ($\mathcal{D}_{23-24}$). The sequence identity between the two sets is calculated and used for further analysis. The structures are divided into two groups based on a sequence identity cutoff of 50.00%, as presented in the main text. A large part of the structure 7wkp_B is unresolved so the RMSD corresponding to it is only for the resolved part. See supplementary table 3 for results of each structure.

Command Used for the dataset upto 2022 ($\mathcal{D}_{22}$):

```
export DATA_PATH=/path/to/data # the dataset is saved here
nucleoseeker \
    --dataset_name name \
    --rfam_cm_path your/rfam/path \
    --exptl_method "X-RAY DIFFRACTION" \
    --resolution 5.0 \
    --year_range 2022 \
    --save 1
```

Command used for the dataset from 2023 to 2024 ($\mathcal{D}_{23-24}$):

```
export DATA_PATH=/path/to/data # the dataset is saved here
nucleoseeker \
    --dataset_name name \
    --rfam_cm_path your/rfam/path \
    --exptl_method "X-RAY DIFFRACTION" \
    --resolution 5.0 \
    --year_range 2023 2024 \
    --save 1
```

##### 3.3 More options for using NucleoSeeker

As in the previous command for Alphafold3 analysis, data for a given `year_range` can be requested by first specifying the start and then teh end year. In addition, some parameters in NucleoSeeker can take multiple arguments, for example, you can provide different values for `exptl_method`, `selected_polymer_entity_types` etc. For a list of possible arguments see table 1

The `exptl_method`, `SOLUTION NMR` can't be used alone, because structures resolved using this technique do not have a `resolution` attribute, thus a dataset will not be created in this case. However, if this is used along with a method which has `resolution` then structures from the other method will be used for creating the dataset.

All the arguments are case-sensitive, and spellings should be correct, otherwise the tool raises errors.

```
export DATA_PATH=/path/to/data # the dataset is saved here
nucleoseeker \
    --dataset_name name \
    --rfam_cm_path your/rfam/path \
    --exptl_method "X-RAY DIFFRACTION" "SOLUTION NMR" \
    --selected_polymer_entity_types "Protein/NA" "Other" \
    --pdbx_keywords "RNA" "RIBOZYME" \
    --resolution 5.0 \
    --year_range 2010 2020 \
    --save 1
```

### 4 Supplementary Tables

Table 1: Arguments for parameters in NucleoSeeker. **NOTE:** The arguments for `Experimental Method`, `Selected Polymer Entity Types`, `PDBx Keywords`, `Polymer Type` are not exhaustive and can change with time. Given arguments should be sufficient for most common use cases. In this list, multiple parameters use the word `Polymer`. The first one is for the five categories of macromolecules in which each structure in the RCSB PDB database is classified. Each structure is composed of multiple entities, the second one is used to filter structures on this classification. The third one checks individual chains in each structure for their type of polymer.

| **Parameter** | **Possible Arguments** | **Usage** |
|---|---|---|
| Structure Determination Methodology | [‘experimental’, ‘computational’] | Single Argument Only |
| RCSB Polymer Entity Type | [‘Protein’, ‘DNA’, ‘RNA’, ‘NA-hybrid’, ‘Other’] | Multiple Arguments Allowed |
| Experimental Method | [‘SOLUTION NMR’, ‘X-RAY DIFFRACTION’, ‘ELECTRON MICROSCOPY’, ‘FIBER DIFFRACTION’, ‘FLUORESCENCE TRANSFER’, ‘SOLID-STATE NMR’] | Multiple Arguments Allowed |
| Selected Polymer Entity Types | [‘Nucleic acid (only)’, ‘Other’, ‘Protein/NA’] | Multiple Arguments Allowed |
| PDBx Keywords | [‘RNA’, ‘DNA/RNA’, ‘RIBOSOME’, ‘RIBOZYME’] | Multiple Arguments Allowed |
| Polymer Type | [‘polyribonucleotide’, ‘polypeptide’] | Single Argument Only |
| Alignment Tool | [‘emboss’, ‘clustal’] | Single Argument Only |

Table 2: Top-L Precision values as obtained from Barnacle and PyDCA. Structure `1duh` failed in both cases because no useful contact map was generated for our criteria and `7vnv` failed because of non-standard residues in the sequence. All others (`1nbs`, `1hr2`, `2il9`, `5ml7`, `8uiw` and `5npm`) failed in Barnacle because it currently doesn't support working with unresolved sections of the structure.

| Family_ID | RCSB_ID | Chain_ID | Barnacle_Top-L Precision | PyDCA_Top-L Precision |
|---|---|---|---|---|
| RF00169 | 1duh | A | nan | nan |
| RF00061 | 1hr2 | A | nan | 0.503 |
| RF00028 | 1kh6 | A | 0.3228 | 0.152 |
| RF00029 | 1kxk | A | 0.8421 | 0.4 |
| RF00011 | 1nbs | A | nan | 0.613 |
| RF00010 | 1u9s | A | 0.1491 | 0.761 |
| RF00164 | 1xjr | A | 0.7857 | 0.468 |
| RF00028 | 1y0q | A | 0.4740 | 0.594 |
| RF01857 | 1z43 | A | 0.8713 | 0.495 |
| RF00234 | 2gcs | B | 0.1920 | 0.52 |
| RF00458 | 2il9 | A | nan | 0.286 |
| RF00521 | 2qwy | A | 0.4815 | 0.423 |
| RF00059 | 3d2v | A | 0.7031 | 0.649 |
| RF00168 | 3dil | A | 0.8851 | 0.698 |
| RF01767 | 3e5c | A | 0.7308 | 0.396 |
| RF01057 | 3npq | A | 0.3500 | 0.333 |
| RF00504 | 3oww | A | 0.7955 | 0.617 |
| RF00380 | 3pdr | X | 0.8519 | 0.534 |
| RF01786 | 3q3z | V | 0.8400 | 0.528 |
| RF00044 | 3r4f | A | 1.0000 | 0.348 |
| RF01852 | 3rg5 | A | 0.8721 | 0.581 |
| RF01510 | 3ski | A | 1.0000 | 0.286 |
| RF01831 | 3suh | X | 0.7228 | 0.505 |
| RF00163 | 3zp8 | A | 0.2500 | 0.111 |
| RF01734 | 4enc | A | 0.7917 | 0.462 |
| RF02001 | 4faw | A | 0.0111 | 0.018 |
| RF00167 | 4fe5 | B | 0.8209 | 0.731 |
| RF01689 | 4frg | B | 0.7143 | 0.333 |
| RF00174 | 4gxy | A | 0.0523 | 0.669 |
| RF01054 | 4jf2 | A | 0.9024 | 0.213 |
| RF00162 | 4kqy | A | 0.7193 | 0.580 |
| RF01725 | 4l81 | A | 0.7634 | 0.479 |
| RF01831 | 4lvw | A | 0.8652 | 0.596 |
| RF03160 | 4oji | A | 0.3214 | 0.617 |
| RF00233 | 4p5j | A | 0.1647 | 0.417 |
| RF01415 | 4pqv | A | 0.4706 | 0.279 |
| RF00379 | 4qln | A | 0.0800 | 0.543 |
| RF02683 | 4rum | A | 0.7831 | 0.484 |
| RF00167 | 4tzx | X | 0.8592 | 0.718 |
| RF01854 | 4wfl | A | 0.7308 | 0.477 |
| RF01750 | 4xwf | A | 0.7115 | 0.438 |
| RF00080 | 4y1m | B | 0.6744 | 0.336 |
| RF03167 | 4yaz | R | 0.7556 | 0.536 |
| RF01750 | 5btp | B | 0.1515 | 0.509 |
| RF03160 | 5dun | A | 0.6667 | 0.635 |
| RF00162 | 5fjc | A | 0.7273 | 0.533 |
| RF00100 | 5lyu | A | 0.5000 | 0.527 |
| RF02540 | 5ml7 | A | nan | 0.286 |
| RF01763 | 5nwq | A | 0.7826 | 0.571 |
| RF00442 | 5t83 | A | 0.0000 | 0.425 |
| RF00442 | 5u3g | B | 0.7600 | 0.442 |
| RF00080 | 6cb3 | A | 0.4752 | 0.320 |
| RF00442 | 6ck5 | A | 0.5862 | 0.349 |
| RF02553 | 6cu1 | A | 0.5455 | 0.423 |
| RF01826 | 6fz0 | A | 0.7500 | 0.421 |
| RF01807 | 6gyv | A | 0.5200 | 0.093 |
| RF02678 | 6jq5 | A | 0.7180 | 0.354 |
| RF02885 | 6lax | A | 0.5000 | 0.164 |
| RF00080 | 6n2v | A | 0.6154 | 0.354 |
| RF00379 | 6n5o | A | 0.6746 | 0.504 |
| RF00102 | 6ol3 | C | 0.8429 | 0.375 |
| RF03165 | 6p2h | A | 0.7826 | 0.450 |
| RF01704 | 6qn3 | A | 0.7600 | 0.320 |
| RF02679 | 6r47 | A | 0.7632 | 0.327 |
| RF03013 | 6tf0 | A | 0.8077 | 0.412 |
| RF02679 | 6ufj | A | 0.7209 | 0.292 |
| RF00174 | 6vmy | A | 0.5000 | 0.264 |
| RF00050 | 6wjr | X | 0.7544 | 0.491 |
| RF02680 | 6xkn | A | 0.0952 | 0.384 |
| RF03013 | 7d7v | A | 0.7719 | 0.333 |
| RF03013 | 7d81 | A | 0.8200 | 0.460 |
| RF03054 | 7elq | A | 0.8000 | 0.489 |
| RF04222 | 7jju | A | 0.5098 | 0.429 |
| RF00525 | 7kga | A | 0.7857 | 0.322 |
| RF00507 | 7mky | A | 0.7727 | 0.258 |
| RF00442 | 7mlw | F | 0.1538 | 0.453 |
| RF00622 | 7qr3 | C | 0.8571 | 0.203 |
| RF02977 | 7wii | V | 0.8000 | 0.38 |
| RF01764 | 8uiw | N | nan | 0.543 |
| RF00036 | 8uo6 | A | 0.0000 | 0.127 |
| RF02541 | 5npm | B | nan | 0.475 |
| RF02541 | 6prv | A | 0.0000 | 0.509 |
| RF00005 | 1ehz | A | 0.8684 | 0.586 |
| RF00005 | 1vtq | A | 0.7467 | 0.646 |
| RF00005 | 3cw5 | A | 0.7922 | 0.652 |
| RF00005 | 6pmo | B | 0.1200 | 0.681 |
| RF00005 | 6ugg | A | 0.8052 | 0.662 |
| RF00005 | 7vnv | A | nan | nan |

### 5 Supplementary Figures

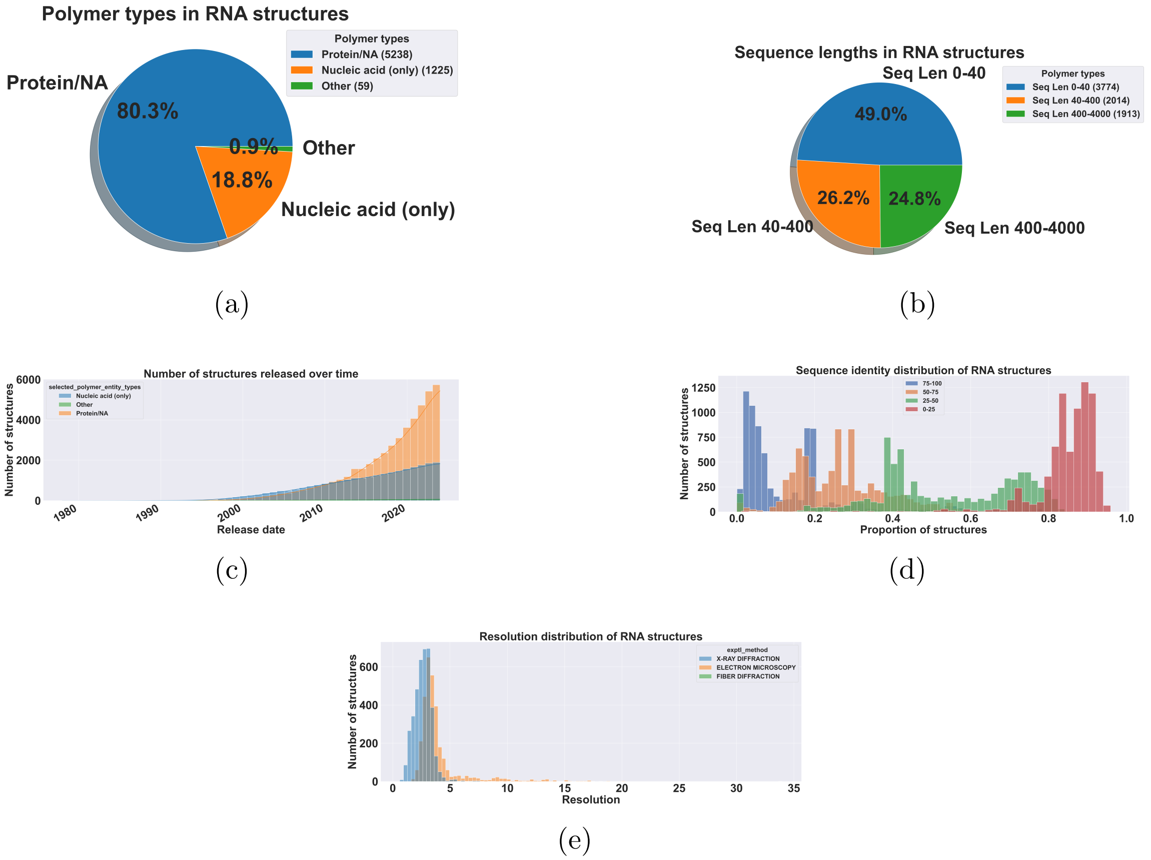

Figure 1: We analysed all the 7704 RNA structures in the PDB database (as of July 2024). They were obtained without applying any filters in NucleoSeeker. (a) We see the composition of different polymer types, most structures are complexes instead of RNA only. (b) Distribution of sequence lengths in RNA structures where it is shown that a large proportion of structures have very short lengths (0-40 residues) (c) Over the years, the RNA structures released at PDB are mostly composed of `Protein` and `RNA` complexes. (d) A large proportion of RNA structures (∼25%) have more than 75% sequence identity. These are some limitations of all the RNA structures which should be closely considered when training a machine-learning model on such a dataset because this redundancy can have a strong impact on performance.

Table 3: Alphafold 3 analysis results for structures divided into two groups based on sequence identity. For this, the dataset was generated using methods described in section 3.2. textttnan values represent the failed predictions. The `Score` here is the `pTM score` provided by Alphafold3. It is described here as the predicted template modeling (`pTM`) score, values higher than 0.8 represent confident high-quality predictions, while values below 0.6 suggest likely a failed prediction. ipTM values between 0.6 and 0.8 are a gray zone where predictions could be correct or incorrect.

| RCSB_ID | RMSD (Å) | Score | Seq Len | Seq Id(%) |
|---|---|---|---|---|
| 7u4a_A | 25.558 | 0.23 | 72 | 52.94 |
| 8c3a_1 | 5.651 | 0.71 | 3359 | 100.00 |
| 8f4o_A | 10.433 | 0.55 | 83 | 57.14 |
| 8fc6_1A | 2.790 | 0.82 | 2915 | 90.00 |
| 8ffr_W | 64.253 | 0.11 | 99 | 55.55 |
| 8fom_QA | 1.209 | 0.8 | 1522 | 100.00 |
| 8fw4_Z | 2.422 | 0.66 | 210 | 50.90 |
| 8g9z_E | 3.434 | 0.52 | 90 | 98.83 |
| 8hnt_B | 28.505 | 0.26 | 128 | 71.09 |
| 8kal_A | 18.417 | 0.24 | 98 | 77.77 |
| 8olv_A | 1.662 | 0.87 | 394 | 100.00 |
| 8pot_B | 2.597 | 0.56 | 87 | 100.00 |
| 8uo6_A | 9.427 | 0.4 | 134 | 72.72 |
| 8axf_Q | 16.901 | 0.3 | 42 | 100.00 |
| 8its_A | 11.548 | 0.3 | 46 | 54.34 |
| 7wii_A | 7.673 | 0.3 | 50 | 42.00 |
| 7wkp_B | 5.715 | 0.25 | 60 | 43.33 |
| 8dp3_R | 20.89 | 0.39 | 90 | 41.67 |
| 8gxb_A | 6.931 | 0.32 | 61 | 41.81 |
| 8hzl_A | 30.245 | 0.29 | 84 | 41.86 |
| 8sh5_R | 6.183 | 0.34 | 88 | 37.78 |
| 8t29_R | 11.482 | 0.29 | 90 | 42.22 |
| 8uiw_N | 12.455 | 0.38 | 128 | 41.17 |
| 9bun_A | 17.009 | 0.28 | 48 | 45.83 |
| 8eyu_B | 14.673 | 0.29 | 49 | 44.89 |
| 8idf_B | 12.052 | 0.32 | 53 | 40.00 |
| 8joz_B | 3.060 | 0.51 | 88 | 47.67 |

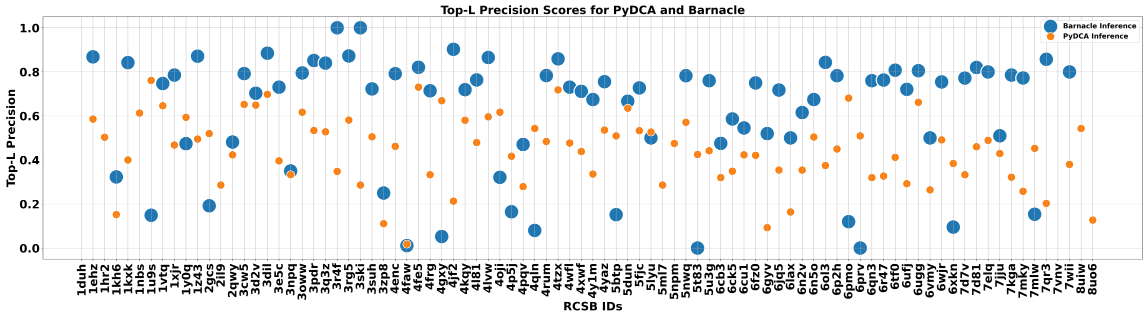

Figure 2: Comparison of Top-L precision scores for PyDCA and Barnacle across various structures of dataset $\mathcal{D}_C$. The x-axis shows RCSB IDs for different protein structures, while the y-axis represents the Top-L precision scores ranging from 0 to 1. Blue circles indicate Barnacle inference scores and orange squares represent PyDCA inference scores. The plot demonstrates the relative performance of these two methods in predicting protein contact maps, with Barnacle generally achieving higher precision scores for most structures. This analysis highlights Barnacle's credibility in contact map prediction and showcases the utility of NucleoSeeker for generating diverse, non-redundant datasets. The dataset for this analysis was obtained using the command specified in section 3.1.

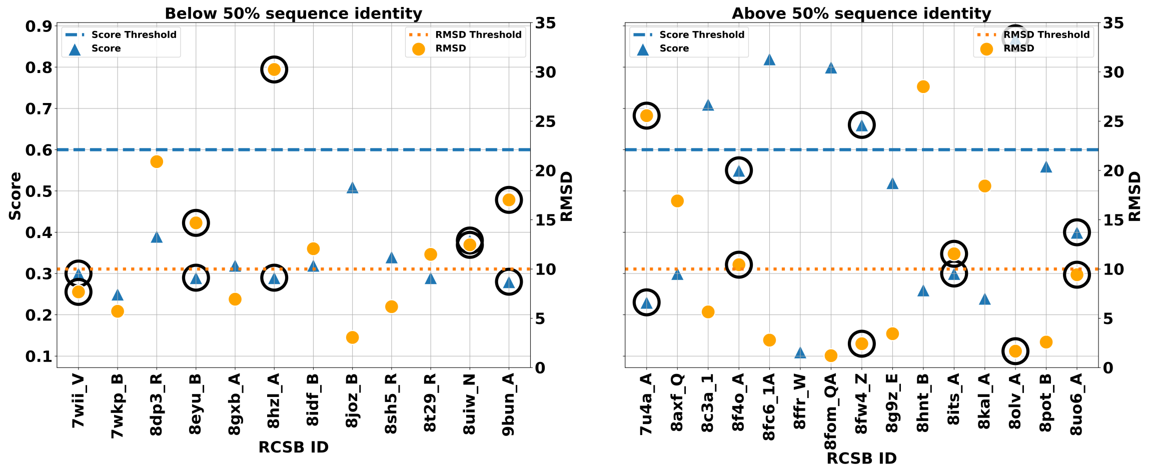

Figure 3: Performance comparison of AlphaFold3 on an unseen dataset (`DATA_23_24`), classified by sequence identity to data (`DATA_22`). The plot is divided into two panels: sequences with below 50% identity (left) and above 50% identity (right) to the training set. Each point represents a structure from the RCSB database, identified by its ID on the x-axis. Two metrics are displayed: **Score (blue triangles)** and **RMSD (orange circles)**, with their respective thresholds shown as dashed lines. The y-axis on the left corresponds to Score values (0-1 scale), while the right y-axis shows RMSD values (0-35Åscale). Circled data points are RNA-only structures. This plot demonstrates how sequence similarity to the training data can influence AlphaFold3's predictive performance, with generally improved scores and lower RMSD values observed for sequences with higher sequence identity to the `DATA_22`. The dataset was generated using the commands given in section 3.2.
